# DTA expression in mTas1r3+ cells lead to male infertility

**DOI:** 10.1101/2020.04.15.042614

**Authors:** Feng Li, Bowen Niu, Lingling Liu

## Abstract

TAS1R taste receptors and their associated heterotrimeric G protein gustducin are strongly expressed in testis and sperm, but their functions and distribution in these tissues were unknown. Using transgenic mouse models, we show that taste signal transduction cascades (mTas1r3-Gnat3-Trmp5) are observed in testis form GFP transgenic mice. It is mTas1rs and mTas2rs, not Gnat3, that was expressed in leydig and sertoli cells. The pattern of mTas1r3 expression was different from that of mTas2r105 expression in seminiferous epithelium. Analysis of the seminiferous epithelium cycle show that both mTas1r3 and mTas2r105 is expressed in the spermatid stage, but mTas2r5 expression is found in spermatocyte stage. Conditional deletion of mTas1r3+ cells leads to male infertility, but do not affect the expression of taste signal transduction cascade during the spermatogenesis. The current results indicate a critical role for mTas1r3+ cell in sperm development and maturation.

## Introduction

The Tas1rs are dimeric Class III GPCRs, with large N-terminal ligand binding domains. Three different Tas1rs have been reported, including Tas1r1, Tas1r2 and Tas1r3. So far, it is always believed that these receptors have the biological function in vivo only as heterodimers, with Tas1r3 serving as an obligate partner for both the umami receptor (Tas1r1+Tas1r3) and the sweet receptor (Tas1r2+Tas1r3) (Nelson et al., 2002; Nelson et al., 2001). Furthermore, knockouts of those genes have been used to identify the role of Tas1rs in taste transduction (Damak et al., 2003). On the other hand, the production of several lines of transgenic mice expressing green fluorescent protein (GFP) from their promoters, particularly Gnat3, Trmp5 and Tas1r3, has revealed an extensive expression in the airways, from the upper airways to the lungs. Furthermore, Tas1rs signaling has been most extensively studied in indigestive system. The sweet receptor is selectively expressed in enteroendocrine L cells, which can release GLP-1 after sugar ingestion, and GLP-1 in turn augments insulin release from the pancreas (Margolskee et al., 2007). The expression of the Tas1rs and their associated G-protein genes has also been reported in mammalian brain, indicating that the Tas1r2/Tas1r3 is a candidate membrane-bound brain glucosensor (Ren et al., 2009; Voigt et al., 2015).

Recently, several works reveal the expression of taste receptors in male reproductive system. Voigt et al (Voigt et al., 2012) replaced the mTas1r1 and mTas2r131 open reading frames with the expression cassettes containing the fluorescent proteins mCherry or hrGFP, respectively. With these transgenic mice, Tas1r1 expression was observed in testis, epididymis mature spermatids. Meanwhile, Tas1r1 expression was also detected in earlier developmental stages of spermatogenesis and Sertoli cells. On the other hand, mTas2r131 expression was observed in spermatocyte and spermatid. In taste cells, the expression of mTas1r1 and mTas2r131 is segregated, whereas co-located in male germ cell. In mature spermatozoa, the expression of the mTas1r1 and mTas2r131 is restricted in distinct segments of the flagellum and the acrosomal cap (Meyer et al., 2012). In situ hybridization and real-time PCR revealed the expression of 35 Tas2rs in testis (Xu et al., 2012). With the humanized mTas1r3 transgenic mice, the genetic deletion of mTas1r3 gene and Gnat3 gene, Mosinger et al (Mosinger et al., 2013) revealed a crucial role for these genes in sperm development and maturation. Interestingly, another study have showed the associations between three single nucleotide polymorphisms (SNPs) in taste receptors genes and male infertility (Gentiluomo et al., 2017).

In previous study, we generated two transgenic mice, one expressing Cre/GFP fusion protein from mTas2r105 promoters (Li and Zhou, 2012), and another expressing Cre/GFP fusion protein from mTas1r3 promoters (Li, 2013). The previous study with the mTas2r105-Cre/GFP mouse revealed mTas2r105 expression in testis (Li and Zhou, 2012). We further provide a hypothesis that taste receptors may be involved in the regulation of spermatogenesis (Li, 2013). Here, we analyse the mTas1r3 expression in testis with the mTas1r3-KO and mTas1r3-Cre/GFP mice. The current results show that the genetic modification of mT1R3 or ablation of Tas1r3+ cells do not affect the expression of taste signal transduction cascade during the spermatogenesis. DTA expression in Tas1r3+ cells results in male infertility.

## Materials and Methods

### BAC-Tas1r3-Cre/GFP construction and generation of transgenic mouse lines

BAC RP23-242K15 containing the entire Tas1r3 gene was purchased from BACPAC Resources, Oakland, USA. Tas1r3 gene contains six exon, and IRES-Cre/GFP fragment is designed to insert into Exon 6. Two homologous arms of 48 bp from the Tas1r3 gene were synthesised into both sides of the primers used to amplify the IRES-Cre/GFP-frt-KanR-frt cassette in the pICGN21 plasmid (forward primer, 5’-AGATGAGAACAGTGGCGGTGGTGAGGCAGCTCAGGGACACAATGAATGAGCCAAGCTATCGAATTCCGCC-3’; reverse primer, 5’-TCGGGGCTTGTTGGCTTAGGAGATTTCTGGTAGGGCTAGGTTCCCTAAA GGG CTATTCCAGAAGTAGTGAGGA-3’). The IRES-Cre/GFP-frt-KanR-frt cassette was introduced into the mTas1r3 gene downstream of the stop codon and upstream of the Poly(A) signal, by means of Red-mediated homologous recombination (Lee et al., 2001), followed by flp-mediated removal of the KanR selectable marker from the BAC-Tas1r3-IRESCre/GFP construct. Restriction digestion with pulsed-field gel electrophoresis and sequencing were used to confirm the correct insertion of the IRES-Cre/GFP gene into Tas1r3 gene.

The BAC-Tas1r3-IRES-Cre/GFP construct was purified and microinjected into the pronuclei of B6 (C57BL/6J) mouse zygotes. The three transgenic founder mice and their progeny were identified using polymerase chain reaction (PCR) with Cre/GFP-specific primers and Tas1r3-specific primers. The expected PCR products were 421 bp with a pair of primers (Forward primer 5’-GATACCTGGCCTGGTCTGGA-3’; Reverse primer 5’-TTTGGGGAAGGGGTCACTGT-3’).

Genotype analyses including R26:lacZbpA^flox^ diphtheria toxin A (DTA) line (Brockschnieder et al., 2006), Tas1r3-KO (Damak et al., 2003), Gnat3-GFP mouse (Wong et al., 1999), Trmp5-GFP (Clapp et al., 2006) were carried out by PCR as previously described. All animal procedures were approved by the committee of Shanghai Public Health Clinical Center and conformed to the International Guiding Principles for Biomedical Research Involving Animals as promulgated by the Society for the Study of Reproduction.

### Primary Sertoli and Leydig Cell Isolation

Primary Sertoli and Leydig cells were isolated by Percoll density gradient centrifugation, as previously described (Chang et al., 2011). Testes from 6- to 8-week-old mice were collected in Enriched DMEM:F12 (Mediatech, Manassas, VA, USA) and placed on ice. After removal of the testes’tunica albuginea, seminiferous tubules were dissociated and transferred immediately into 10 mL Enzymatic Solution 1 (Enriched DMEM:F12, 200 μg/mL deoxyribonuclease I, 0.5 mg/mL collagenase IA). Tubules were incubated for 15–20 min at 35°C in a shaking water bath at 80 oscillations (osc)/min, then layered over 40 mL 5% Percoll (Sigma-Aldrich, St. Louis, MO, USA)/95% 1×Hank’s balanced salt solution (HBSS; Invitrogen, Carlsbad, CA, USA) in a 50-mL conical tube and allowed to settle for 20 min. Leydig cells were isolated from the top 35 mL Percoll previously described (Chang et al., 2011). The bottom 5 mL Percoll were transferred to a fresh 50-mL conical tube containing 10 mL Enzymatic Solution 2 (Enriched DMEM:F12, 200μ g/mL deoxyribonuclease I, 1 mg/mL trypsin). Tubules were incubated for 20 min at 35°C and 80 osc/min. After incubation, 3 mL charcoal-stripped FBS (Thermo Fisher Scientific, Waltham, MA, USA) were immediately added to halt the digestion. The digested product was filtered through a 70-μm cell strainer and again through a 40-μm cell strainer. Sertoli cells were isolated from cells retained by the filters previously described (Chang et al., 2011).

### RNA Isolation, cDNA Synthesis and RT-PCR

RNA was isolated from primary sertoli and leydig Cell using Tri-Reagent (Ambion) following the manufacturer’s protocol. Purification and reverse transcription RNA with TransScript One-Step gDNA Removal and cDNA Synthesis SuperMix (TransGen Biotech), PCR amplification with TransTaq DNA Polymerase High Fidelity (TransGen Biotech), all the steps are according to manufacturer’s protocol. Sequences of primers used and the predicted amplification sizes are as followed: Tas2r105 Forward 5’-GGC ATC CTC CTT TCC ATT-3’, Reverse 5’-ACC GTC CTT CAT CAC CTT C-3’ 447bp; Tas2r106 Forward 5’-TCA CAG GCT TGG CTA TTT-3’, Reverse 5’-TTG AGA AGA ATG TGG CTT AC-3’, 397bp; Tas2r113 Forward 5’-TAC CCA GCA TTA CAC GAAA-3’, Reverse 5’-CAG GGA GAA GAT GAG CAA A-3’ 402bp; Tas2r134 GAA TAG CCG TCC TAA ACA A-3’, Reverse 5’-TCC AGA TGC CGA TAC AGT-3’ 433bp; Tas2r143 Forward 5’-GAG GAT TTC CCA GTT AGT TC-3’, Reverse 5’-ATG GTA TGT GCC TGA GTA TG-3’ 324bp; Gnat3 Forward 5’-ATG GCT ACA CTG GGG ATT-3’, Reverse 5’-TTC TGT TCA CCT CCT CAT CT-3’ 52 °C 466bp.

### Histology and immunostaining procedure

For immunocytochemistry, mice were perfused transcardially with 2–4% paraformaldehyde (PFA) in phosphate-buffered saline (PBS; pH 7.2–7.4). The testis were dissected, post-fixed in PFA for 2–12 hours and cryoprotected in 30% sucrose in PBS at 4°C overnight. After sectioning on a cryostat, 10–12 µm sections were collected onto Superfrost Plus Microscope slides (Fisher Scientific, Loughborough, UK). The polyclonal primary antibodies used were specific for GFP (goat ab-5450, rabbit ab-6556; Abcam, Cambridge, UK), PLC-β2 (rabbit sc-206; Santa Cruz Biotechnology, Santa Cruz, USA), GNAT3 (rabbit sc-395; Santa Cruz Biotechnology), Tas1r1 (rabbit sc-50308; Santa Cruz Biotechnology, Santa Cruz, USA), Tas1r2 (rabbit sc-50306; Santa Cruz Biotechnology, Santa Cruz, USA), Tas1r3 (rabbit sc-22459; Santa Cruz Biotechnology, Santa Cruz, USA). The rabbit polyclonal antibody against Trpm5 (1028–1049 amino acids) was as described by Perez (Perez et al., 2002).

Staining against Tas1r1, Tas1r2, Tas1r3, GNAT3 and PLC-β2 was performed using the standard immunocytochemical procedure. Cryosections were washed in PBS (3 × 10 min), placed into blocking solution [1% bovine serum albumin (BSA), 1% normal horse serum, and 0.3% Triton X-100 in PBS] for 1–2 hr, and then incubated in a mixture of the polyclonal, primary antisera: rabbit anti-GFP (1:500-1000 dilution) in blocking solution. Primary antibody incubation lasted for 36–48 hr at 4 °C, and then sections were washed in PBS 3 × 10 min and incubated for 2–18 hr in a mixture of secondary antibodies: Alexa568 goat anti-rabbit (1:400; Molecular Probes, USA). The slides then were washed one time for 10 minutes in 0.1 M PB and two times for 10 minutes in 0.1 M PBS before coverslipping slides with Fluormount G (Southern Biotechnology Associates, USA). Fluorescent images were captured using a Leica TCS SP2 Spectral Confocal Microscope (Leica Microsystems Inc., Mannheim, Germany).

Staining against GFP, Tas1r1, Tas1r2, Tas1r3, GNAT3, PLC-β2, and Trpm5 was performed using the standard immunocytochemical procedure according to the manufacturer’s instructions (VECTASTAIN Elite ABC Kit; Vector Laboratories, Burlingame, USA). The standard immunocytochemical procedure used an avidin-biotin-peroxidase complex (ABC; Vector Laboratories, Burlingame, CA) and 3,38-diaminobenzidine (DAB, Sigma). Cryosections were washed in four 4-minute washes of 0.1 M PBS, pH 7.4, containing 0.3% Triton X-100. The slides subsequently were incubated in blocking solution [1% bovine serum albumin (BSA), 1% normal horse serum, and 0.3% Triton X-100 in PBS] for 1–2 hr, followed by a 24 h incubation with the primary antibody at 4°C. The slides were washed with PBS four times followed by a 45-minute application of the biotin-conjugated secondary antibody. Three additional washes in PBS preceded both the 30-minute application of ABC and the 10-minute incubation with a PBS solution containing 0.5 mg/ml DAB, and 0.01% H_2_O_2_ to tint the reaction product blue. Counterstaining is carried out with standard hematoxylin staining and bright field images of the sections were captured digitally.

### Immunostaining blotting

For immunoblotting, immunoprecipitates or protein extracts (50 lg) were resolved by one-dimensional SDS-PAGE (12% acrylamide) under reducing conditions and electrophoretically transferred to polyvinylidene difluoride membranes (GE Amersham). Membranes were probed with primary antibodies against mTas1r1, mTas1r2, mTas1r3, Gnat3 or β-actin (ACTB; 1:10 000 dilution; product sc-47778; Santa Cruz Biotechnology) diluted in Tris-buffered saline with 0.1% tween 20 containing 5% bovine serum albumin (Jackson ImmunoResearch Laboratories) or 5% dried milk. Following incubation with a horseradish peroxidase-conjugated secondary goat anti–rabbit antibody (1:10 000 dilution; product W401B; Promega), the protein bands were visualized by chemiluminescence, using Immobilon Western chemiluminescent horseradish peroxidase substrate (Millipore).

### Sperm count

Cauda epididymides were dissected and minced in phosphate-buffered saline solution. Sperm were squeezed out with fine forceps and allowed to disperse in phosphate-buffered saline at room temperature for 10 min, followed by repeated pipetting. Samples were fixed in 4% paraformaldehyde. Sperm were counted using a hematocytometer according to previous literature (Kovacs and Foote, 1992; Somfai et al., 2002). Sperm counting was performed four times for each sample.

### Evaluation of the percentage of sperm motility and observation of sperm morphology

The percentage of sperm motility after 24-h incubation was tested using grading of motility by WHO criteria with respect to the fertilization of oocytes in vitro (Sukcharoen and Keith, 1996). To observe the mass motility, one drop of semen was placed on a pre-warmed (37°C) slide under microscope at 10x. Mass motility was scored into 0 to 5 scales every two hours according to the protocol suggested by Sukcharoen and Keith (Sukcharoen and Keith, 1996). Routine staining procedure was used to observe sperm morphology (Somfai et al., 2002). Equal drops of trypan blue and diluted semen were mixed on slides with the edge of another slide to make smears. After drying at room temperature (RT), slides were fixed in 4% Paraformaldehyde for 2 min, then stained in 7.5% Giemsa for 12-20 hrs (overnight) at RT. After drying, they were mounted and coverslipped. Slides were evaluated by bright field light microscopy using 100 × oil immersion objectives, as previously described (Kovacs and Foote, 1992; Somfai et al., 2002).

## Results

### Taste signal transduction cascades (mTas1r3-Gnat3-Trmp5) are observed in testis form GFP transgenic mice

Previous study has shown the expression of taste receptors in testis. In order to further verify the expression of taste receptor and signal transduction cascades in the testis, we directly observe the testis from Gnat3-GFP (Fig. 1A), Trmp5-GFP (Figure-1B) and mTas1r3-GFP (Fig. 1C and 1D) transgenic mice with Stereoanatomical fluorescence microscope. More GFP signals was found in the testis from Gnat3-GFP transgenic mice (Fig. 1A), the fewest GFP signals was observed in the testis from mTas1r3-GFP transgenic mice (Fig. 1C and 1D). DTA expression lead to the loss of GFP signals in testis from mTas1r3-Cre/GFP-DTA transgenic mice (Fig. 1E and 1F).

**Figure-1.**
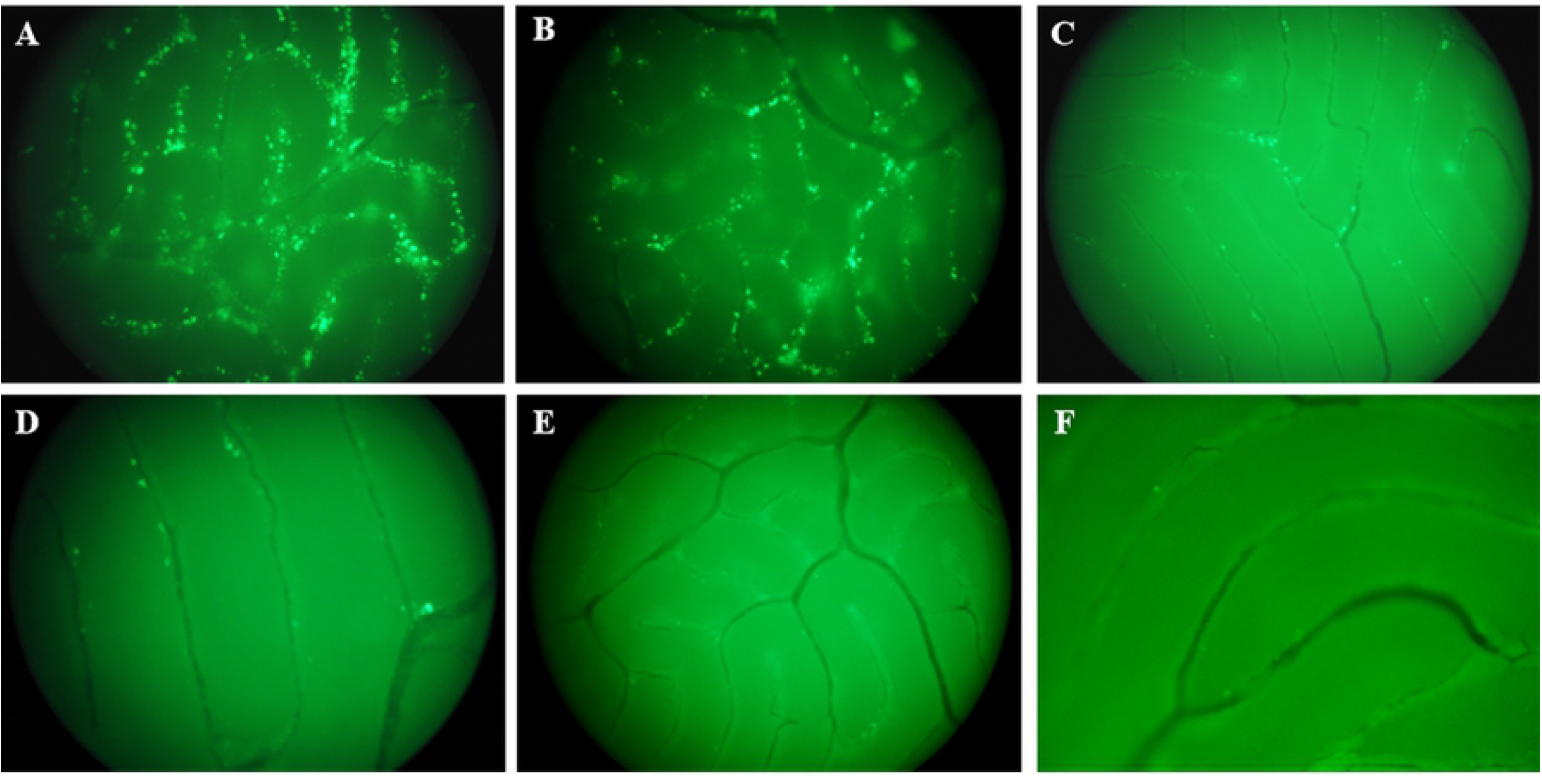
GFP expression in testis from the Gnat3-GFP, Trmp5-GFP and mTas1r3-Cre/GFP transgenic mice. (A) Testis of the Gnat3-GFP transgenic mice. (B) Testis of the Trmp5-GFP transgenic mice. (C-D) Testis of the mTas1r3-Cre/GFP transgenic mice. (E-F) Testis of the mTas1r3-Cre/GFP-DTA transgenic mice.

In order to further check the expression of taste receptor and signal transduction cascades in the seminiferous tubules, immunohistochemistry with anti-GFP was employed to analyze GFP expression in seminiferous tubule from Gnat3-GFP (Fig. 2A-2C), Trmp5-GFP (Fig. 2D-2F) and mTas1r3-Cre/GFP (Fig. 2G-2I) transgenic mice. Immunohistochemistry with anti-GFP revealed the expression of Gnat3 (Figure 2 A-C), Trmp5 (Figure 2 D-F) in later spermatid during the spermatogenesis. Tas1r3 expression was also detected in spermatid and leydig cell (Fig 2G and 2H).

**Figure-2.**
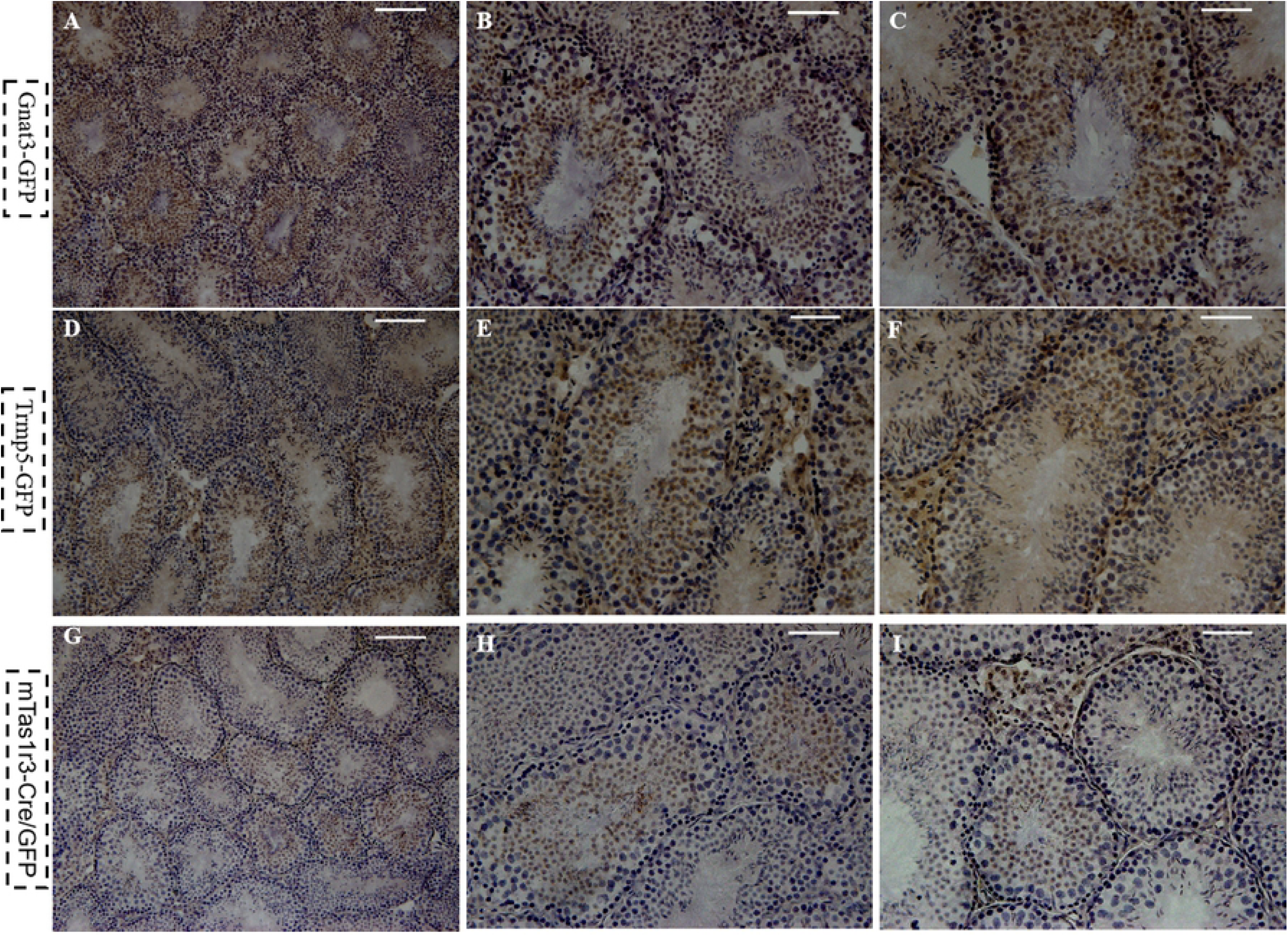
Immunohistochemistry analysis reveals the expression of Gnat3 in Gnat3-GFP transgenic mouse (A-C), Trmp5 in Trmp5-GFP transgenic mouse (D-F) and mTas1r3 in mTas1r3-Cre/GFP transgenic mouse (G-I). (A) Low magnification and (B-C) High magnification in the Gnat3-GFP transgenic mouse. (D) Low magnification and (E-F) High magnification in the Trmp5-GFP transgenic mouse. In the mTas1r3-Cre/GFP transgenic mouse, immunohistochemistry analysis with anti-GFP indicate the GFP expression in spermatogenesis, including leydig cells (G-I). (G) Low magnification. (H-I) High magnification. GFP expression is detected in spermatid phase in the three transgenic mice. Scale bar A D G 100 μm, B C E F H I 50 μm.

### mTas1rs and mTas2rs was expressed in leydig and sertoli cells

We separated primary leydig, sertoli cells and spermatogenic cell, the expression of mTas1r1, mTas1r2 and mTas1r3 was examined by immunostaining blotting in these cells (Fig. 3A). Unexpected, Gnat3 expression was only observed in spermatogenic cells (Fig. 3A). We also analyzed the expression of mTas2rs in testis somatic cells with RT-PCR. Several mTas2rs transcripts was detected in leydig and sertoli cells including mTas2r105, mTas2r106 and mTas2r143 (Fig. 3B). Gnat3 transcripts failed to be detected in leydig and sertoli cells (Fig. 3B). We further employ immunostaining to analyze the expression of mTas1rs in primary leydig and sertoli cells. Confocal analysis showed the expression of mTas1r1, mTas1r2 and mTas1r3 in these cells (Fig. 3C-3J). In short, current results reveals the expression of mTas1rs and mTas2rs in leydig, sertoli cells.

**Figure-3.**
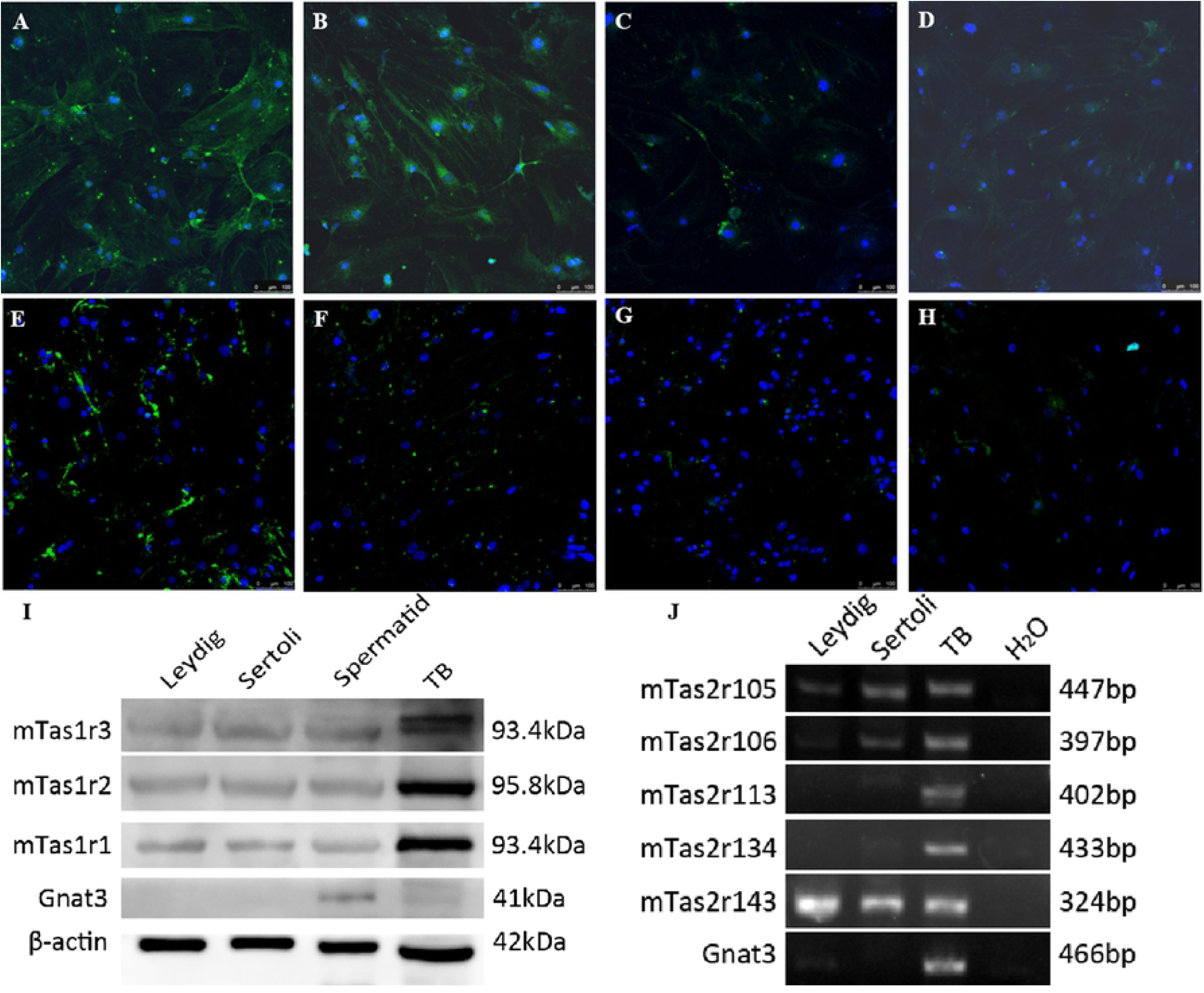
Immunofluoresence staining images for mTas1r3 (A), mTas1r2 (B), mTas1r1 (C) in sertoli cells. Immunofluoresence staining images for mTas1r3 (E), mTas1r2 (F), mTas1r1 (G) in leydig cells. Negtive control in sertoli cells (D) and leydig cells (H). Nuclei were visulaized with DAPI (blue). Scale bar 100 μm. The expression level of mTas1rs in sertoli cells and leydig cells are examined by immunoblotting. GNAT3 is not expressed in sertoli cells and leydig cells (I). The expression of mTas2rs in sertoli cells and leydig cells are examined by RT-PCR. Gnat3 is not expressed in sertoli cells and leydig cells (I).

### mTas1r3 expression was different from mTas2r5 expression in seminiferous epithelium

In order to learn more about the mTas1r3 expression during spermatogenesis, the cycle of the seminiferous epithelium was analyzed in the mTas1r3-GFP transgenic mice. It was found that GFP expression was detected in most of spermatid stage (Fig. 4A-4D, stage 8, 61.5 %; stage 9-12, 28.2 %). GFP expression was also detected in spermatogonium (Fig. 4A and 4D arrow) and spermatocyte (Fig. 4A triangle, 4B 4D star, 10.3 %). In addition, we also observed the cycle of seminiferous epithelium from mTas2r5-Cre/GFP transgenic mice. GFP expression was detected in spermatid (Fig. 5A-5D and 5H, stage 8, 68%) and spermatocyte (Fig. 5A and 5C-5G, 32%). It was noted that GFP expression was observed in pachyene, diplotene spermatocyte, and secondary spermatocyte as well.

**Figure-4.**
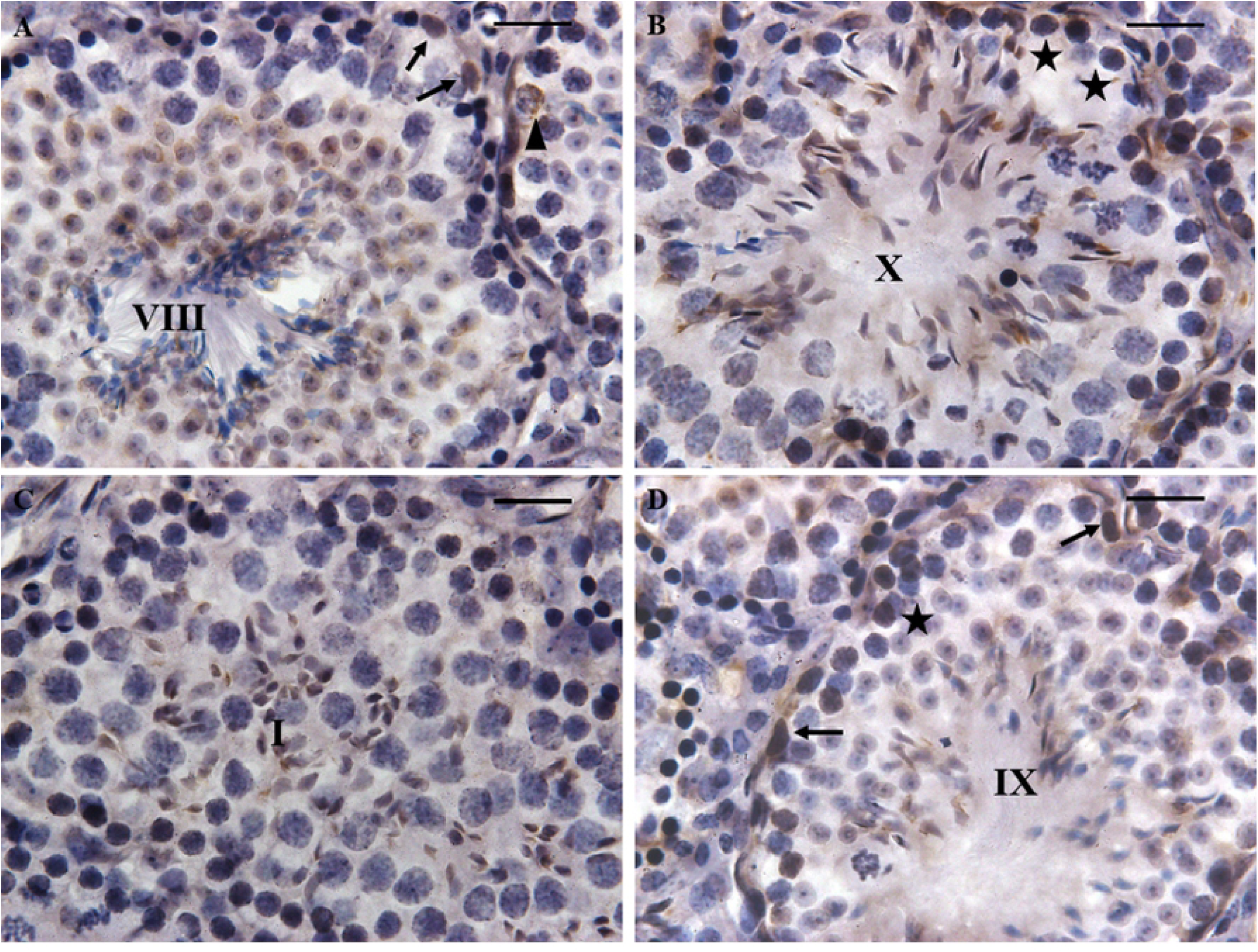
mTas1r3 expression is observed in spermatid. mTas1r3 expression is found in step 8 spermatid of sage VIII (A), step 10 spermatid of stage X (B), step 13 spermatid of stage I (C) and step 11 spermatid of stage IX (D). mTas1r3 expression is also detected in spermatogonium (A D arrow), spermatocyte (A triangle, B D star). Scale bar 20 μm.

**Figure-5.**
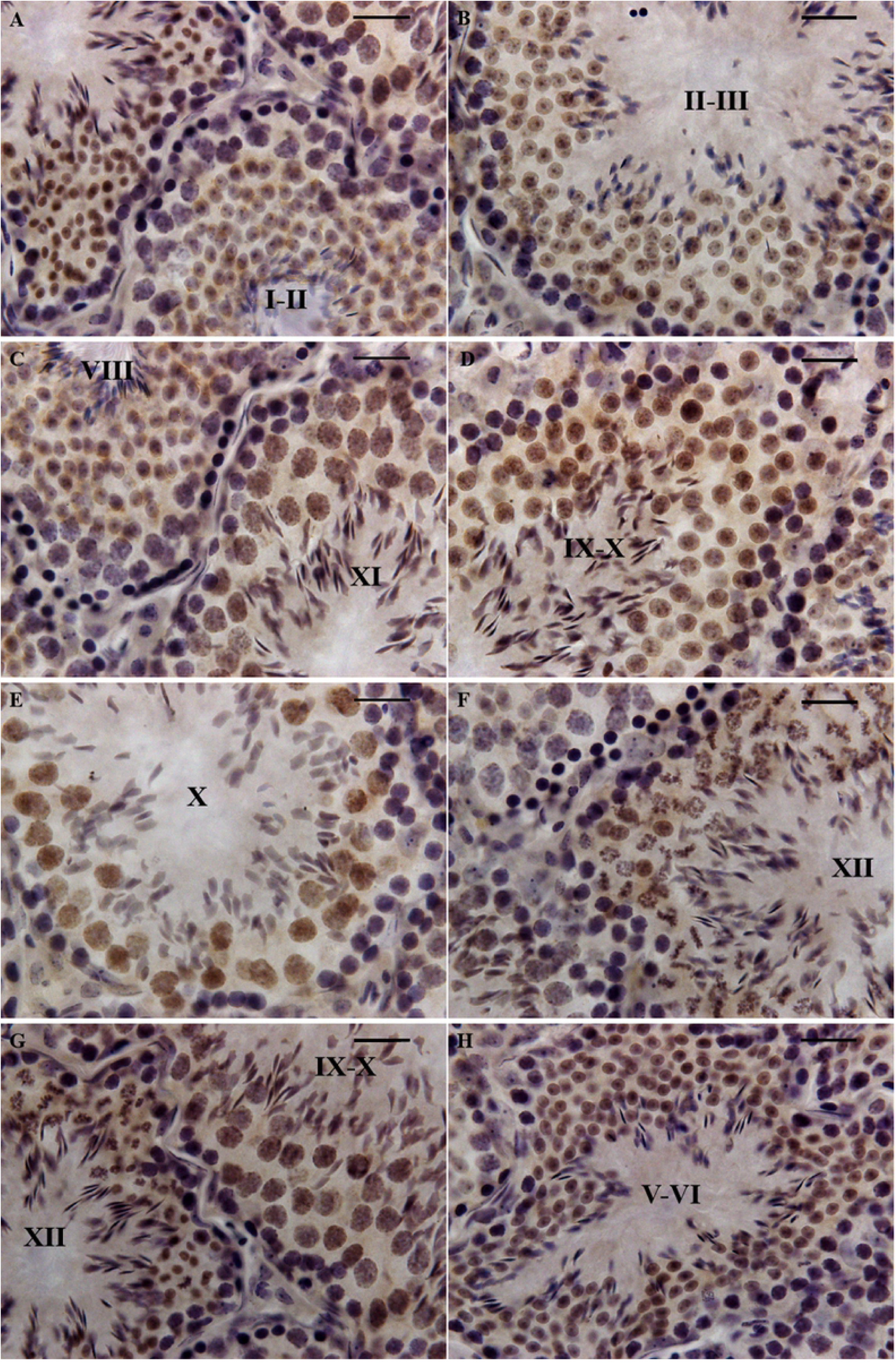
mTas2r105 expression is observed from spermatocyte to spermatid. (A) spermatid in stage I-II. (B) spermatid in stage II-III. (C) spermatid in stage VIII and spermatocyte in stage XI. (D) spermatocyte in stage IX-X. (E) spermatocyte in stage X. (F) meiotic cell division in stage XII. (G) meiotic cell division in stage XII and spermatocyte in stage IX-X. (H) spermatid in stage V-VI. Scale bar 20 μm.

### DTA expression in mTas1r3+ cells decrease sperm mobility

To investigate the function of mTas1r3 during spermatogenesis, the breed data was analysed in mTas1r3-Cre/GFP transgenic mice, mTas1r3-KO mice and mTas1r3-Cre/GFP-DTA. The infertility was observed in male double transgenic mouse (mTas1r3-Cre/GFP-DTA). Additionally, after counting sperm in epididymis, the mTas1r3-Cre/GFP-DTA mice had less sperm concentration than WT males (Fig. 6A). Ratio of testis to body weight was also not significantly different among the group (Fig. 6B). Subsequently, sperm mobility was also checked. Sperm mobility was not significantly different for fresh sperm just removed from epididymis. However, sperm mobility was significantly decreased after cultured for several hours at 37°C (Fig. 6C), indicated that spermatogenesis was affected both in mTas1r3-Cre/GFP-DTA and mTas1r3-KO. Area of semiferous tube from the mTas1r3-Cre/GFP-DTA mice is significantly smaller than that from WT C57BL/6 and mTas1r3-Cre/GFP males (Fig. 6D). About 20% of spermatozoa from the double transgenic mouse (mTas1r3-Cre/GFP-DTA) were abnormal (flipped heads, Head tail separation, flagella with tight loops) (Fig. 6H). These are at least twice as many abnormalities as typically observed in WT C57BL/6 (Fig. 6E) and mTas1r3-Cre/GFP (Fig. 6F) males. The most significantly abnormalities noted were flipped heads (∼8%) and Head without tail (∼8%). In a word, DTA expression in mTas1r3+ cells influence on spermatogenesis with an unknown mechanism.

**Figure-6.**
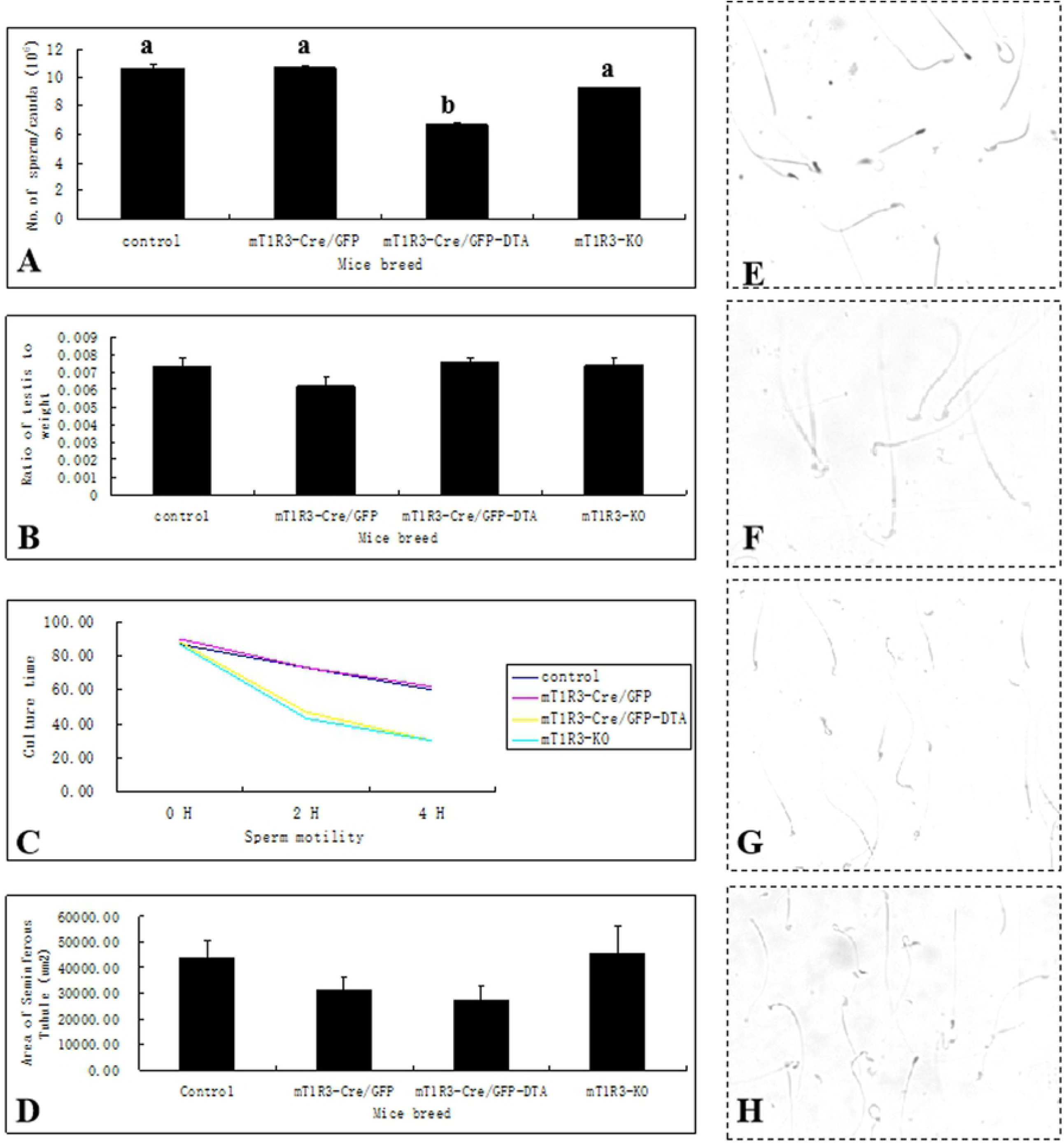
The conditional deletion of mTas1r3+ cells significantly decrease sperm motility. (A) sperm concentration does not differ statistically among the transgenic groups. (B) Ratio of testis to weight is not significantly different among the four groups. (C) Survival curve of sperm in vitro. Sperm motility is significantly decreased for the mTas1r3-KO and mTas1r3-GFP/Cre-DTA mice after 4 H incubation in 37C. (D) Area of seminiferous tubules from mTas1r3-GFP/Cre-DTA mice is significantly decreased. (E) Spermatozoa from control mice. (F) Spermatozoa from the mTas1r3-GFP/Cre transgenic mice. (G) Spermatozoa from the mTas1r3-KO transgenic mice. (H) Abnormal spermatozoa from the mTas1r3-DTA transgenic mice.

### Taste signal transduction cascades (GNAT3-Gγ13-PLCβ2-Trmp5) are still detected in testis from the mT1R3-DTA mouse

The expression of taste signal transduction cascades were further investigated in those transgenic or mTas1r3-KO mouse. As expected, the expression of Gnat3 and PLC-β2 was detected in spermatogenesis in control (Fig. 7A and 7E), mTas1r3-Cre/GFP (Fig. 7B and 7F), mTas1r3-Cre/GFP-DTA (Fig. 7C and 7G) and mTas1r3-KO (Fig. 7D and 7H) mouse.

**Figure-7.**
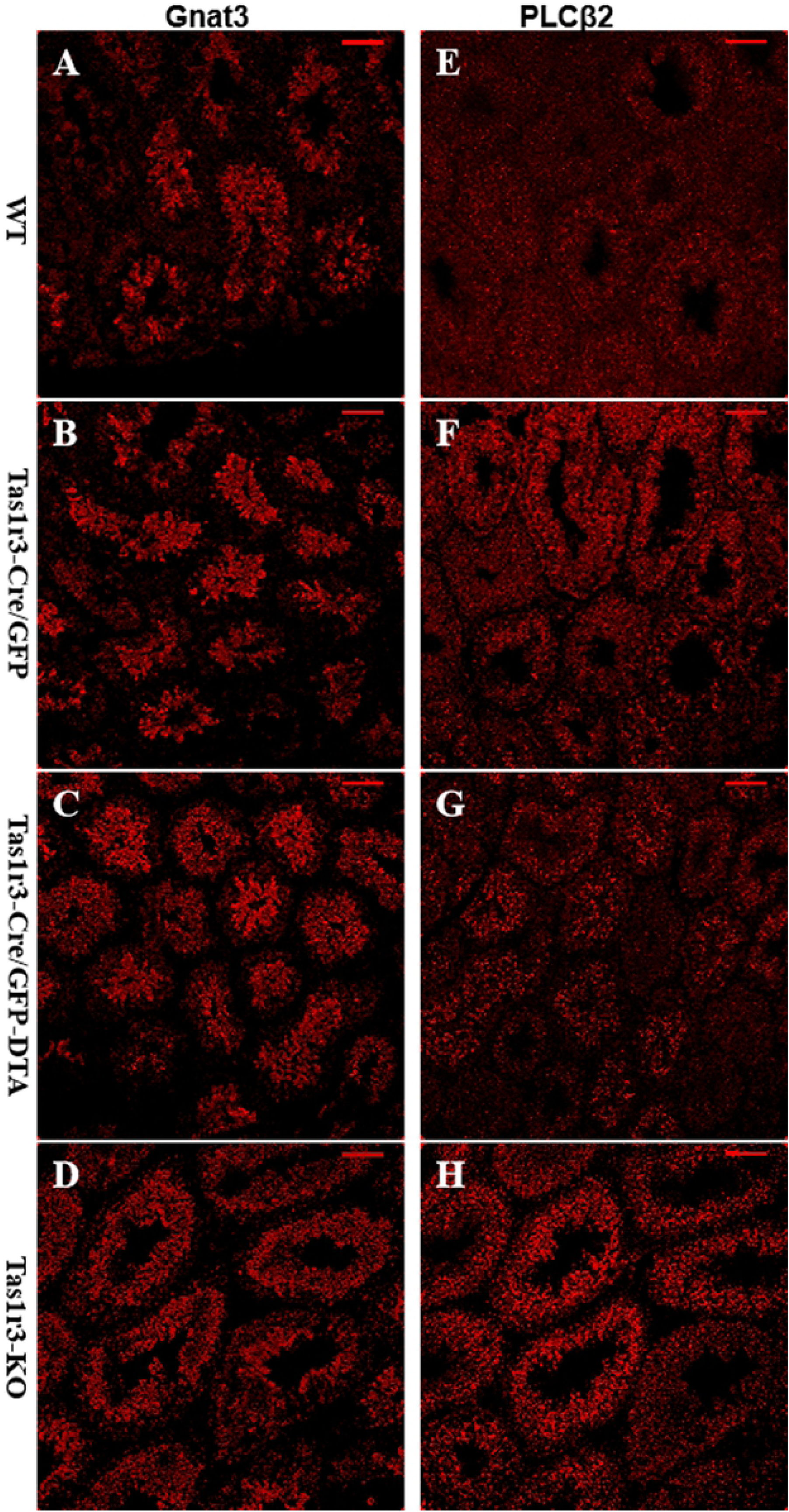
Confocal analysis reveals the expression of taste signal transduction cascades in testis from the transgenic and knockout mouse. (A) Gnat3 expression in control mouse. (B) Gnat3 expression in the mTas1r3-GFP/Cre transgenic mouse. (C) Gnat3 expression in the mTas1r3-GFP/Cre-DTA transgenic mouse. (D) Gnat3 expression in the mTas1r3-KO mouse. (E) PLC-β2 expression in control mouse. (F) PLC-β2 expression in the mTas1r3-GFP/Cre transgenic mouse. (G) PLC-β2 expression in the mTas1r3-GFP/Cre-DTA transgenic mouse. (H) PLC-β2 expression in the mTas1r3-KO mouse. Scale bar A-H 100 μm.

To investigate whether genetic deletion of Tas1r3 or loss of Tas1r3+ cells affected the spermatogenic cycle in mice, Trmp5 expressions were analyzed in adult mice during the 12th spermato-genic cycle. The results indicated that Trmp5 have a unique expression pattern throughout the spermatogenic cycle. Strong Trmp5 immunolabelling was present in the round spermatids at stages VII–VIII and the elongated spermatids at stages 9-15 (Fig. 8A-8F). After loss of mTas1r3+ cells (Fig. 8A1-8F1) or genetic deletion of Tas1r3 (Fig. 8A2-8F2), positive immunostaining was still observed in the round spermatids at stages VII–VIII and the elongated spermatids at stages 9-15.

**Figure-8.**
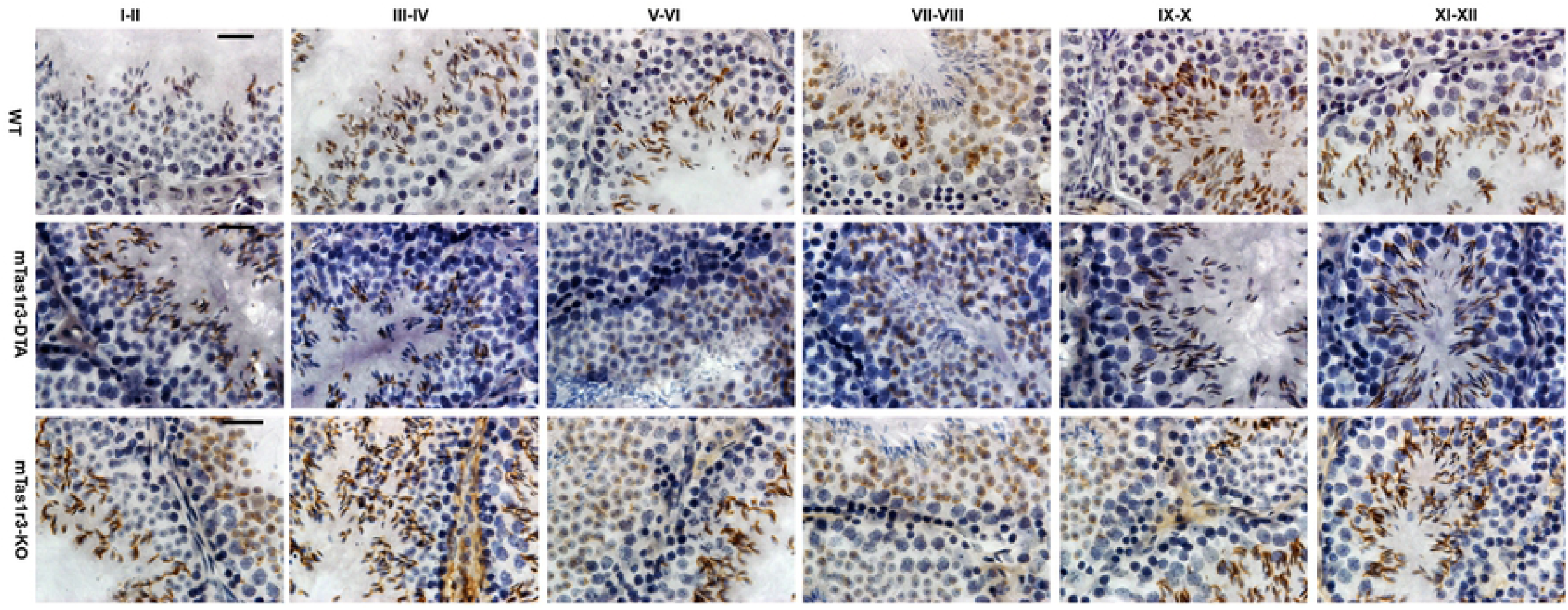
Expression of Trmp5 is observed in the round spermatids at stages VII–VIII and the elongated spermatids at stages 9-15 during the spermatogenic cycle. (A-F) WT control mice. (A1-F1) mTas1r3-GFP/Cre-DTA transgenic mice. (A2-F2) mTas1r3-KO mice.

These intriguing results indicated that genetic deletion of mTas1r3 or loss of Tas1r3+ cells do not affect the expression of taste signalling transduction cascades including Gnat3, PLC-β2 and Trmp5.

## Discussion

We found that Tas1rs are expressed in Leydig and Sertoli cells. We also detected mRNAs for several mTas2rs including mTas2r105 and mTas2r106 in Leydig and Sertoli cells. These data are in agreement with previous reports, which reveal the expression of mTas1rs and mTas2rs in testis (Iwatsuki et al., 2010; Meyer et al., 2012; Voigt et al., 2012; Xu et al., 2012). It is noteworthy that the pattern of mTas2r105 expression differ in striking ways from that of mTas1r3 during spermatogenesis. mTas2r105 expression is widely seen during spermatogenesis including spermatogonial, spermatocyte and spermatid phase. On the other hand, Trmp5 expression is limited to the spermatid phase, and Gnat3 expression is not found in spermatocyte phase and testis somatic cells (leydig and sertoli cells). It thus appears that many taste signaling molecules originally identified in taste cells are expressed also in testis, but they are not as concentrated in a certain type of testis cell as in taste bud. Other signaling molecules may be involved in the downstream of the sweet and bitter taste-sensing receptor in testis. The previous study have shown that the T1R3 receptor is likely coupled with Gas, and mediated the anti-adipogenic signal through activation of Gas in 3T3-L1 cells (Masubuchi et al., 2013).

Agreement with our finding, Tas1r1/Tas1r3 has been found to be present broadly in mouse tissues and many types of cultured cells (Max et al., 2001; Wauson et al., 2012). Even robust expression of Tas1r1/Tas1r3 is found in mice heart (Foster et al., 2013). Further study reveals that amino acids bind Tas1r1/Tas1r3 leading to activation of PLC-β, calcium entry, and ERK1/2, thus stimulating the mammalian target of rapamycin complex 1 (mTORC1) (Wauson et al., 2012). Mammalian mTOR is believed to be a central controller of cell growth, metabolism and aging, coordinating cell growth, protein translation, and autophagy with the availability of nutrients, growth factors, and energy (Dazert and Hall, 2011; Duan et al., 2016; Li and Cheng, 2016; Zoncu et al., 2011). We speculate that Tas1r1/Tas1r3 detects extracellular amino acids, transmits that information to mTORC1 and is involved in regulating cell growth and metabolism in spermatogenesis.

The present study also shows that the pattern of mTas2r105 expression was obviously differ from that of mTas1r3 expression during the spermatogenesis. mTas1r3 expression was mostly observed in mid-later spermatid. In contrast, mTas2r105 expression was observed throughout the spermatogenesis including spermatogonial, spermocyte and spermatid. Thus, DTA expression in mTas2r105+ cells lead to smaller testis (Li, 2013; Li and Zhou, 2012). In taste bud, mTas2r105 is not co-expressed with mTas1r3 (Adler et al., 2000; Chandrashekar et al., 2000). The previous study has also shown that mTas2rs is observed in a subtype of cells distinguished from Tas1r3 positive cells in extra-oral tissue (Clark et al., 2015; Gu et al., 2015; Liu et al., 2015; Voigt et al., 2012). Furthermore, the previous studies have revealed that the expression of mTas1r1 was detected during the earlier developmental stages of spermatogenesis and Sertoli cells (Meyer et al., 2012). The stronger expression of mTas2r131, revealed by hrGFP Fluorescence intensity in mTas2r131^BLiG/BLiG^ mice, was also found in developmental stages of progenitor cells of spermatozoa. Additionally, the expression of mTas1r1 and mTas2r131, separated in taste cells, can partially co-localize in male germ cells (Meyer et al., 2012; Voigt et al., 2012).

Another finding in this study is that DTA expression in mTas1r3+ cells lead to male infertility. Although mTas1r3 expression is mainly observed in mid-later spermatid, loss of mTas1r3 positive cells in testis may be critical during spermatogenesis. Two possibilities should be considered. Firstly, taste genes (such as mTas1r1, mTas1r2, mTas1r3 and their associated G-protein) are detected in mammalian brain, including the paraventricular and arcuate nuclei of the hypothalamus, the CA fields and dentate gyrus of the hippocampus, the habenula, and cortex (Ren et al., 2009). mTas1r3 positive cells may play a role in levels of hypothalamo-pituitary-gonadal feedback loop (Wauson et al., 2012). In addition, the present study demonstrated that Tas1r3 is expressed in leydig and sertoli cells. DTA expression should result in loss of function in mTas1r3 positive cells, which may also contribute to male infertility. Recent study also revealed that polymorphism of several mTas2rs and mTas1r2 was proved to be functional and showed a profound effect on human male infertility (Gentiluomo et al., 2017).

In conclusion, our data suggest a detailed expression pattern of mTas1r3, mTas2r105 and other signal molecules. DTA expression in mTas1r3+ cells finally lead to male-infertile. mTas1r3 and mTas2r105 is expressed in a subtype of cells during spermatogenesis, which is distinct from Gnat3 or Trmp5 positive cells, indicating that there existed a different signaling pathway from taste bud. This unique sweet and bitter-sensing receptor may possibly be a potential target for treatment of male infertility.

## ACKNOWLEDGEMENTS

This work is supported by the Mouse Genome Editing Lab and the followed project: National Science and Technology Major Project (2017ZX10304402-001-006, 2017ZX10304402-001-012); Shanghai professional platform for High-level Biosafety Pathogenic Microorganism Detection (18DZ2293000); the Start-on Funding (KY-GW-2017-06, KY-GW-2018-11 and KY-GW-2019-11) from Shanghai Public Health Clinical Center.

## References

Adler, E., M.A. Hoon, K.L. Mueller, J. Chandrashekar, N.J. Ryba, and C.S. Zuker. 2000. A novel family of mammalian taste receptors. Cell. 100:693–702.

Brockschnieder, D., Y. Pechmann, E. Sonnenberg-Riethmacher, and D. Riethmacher. 2006. An improved mouse line for Cre-induced cell ablation due to diphtheria toxin A, expressed from the Rosa26 locus. genesis. 44:322–327.

Chandrashekar, J., K.L. Mueller, M.A. Hoon, E. Adler, L. Feng, W. Guo, C.S. Zuker, and N.J. Ryba. 2000. T2Rs function as bitter taste receptors. Cell. 100:703–711.

Chang, Y.F., J.S. Lee-Chang, S. Panneerdoss, J.A. MacLean, 2nd, and M.K. Rao. 2011. Isolation of Sertoli, Leydig, and spermatogenic cells from the mouse testis. BioTechniques. 51:341–342, 344.

Clapp, T.R., K.F. Medler, S. Damak, R.F. Margolskee, and S.C. Kinnamon. 2006. Mouse taste cells with G protein-coupled taste receptors lack voltage-gated calcium channels and SNAP-25. BMC Biol. 4:7.

Clark, A.A., C.D. Dotson, A.E. Elson, A. Voigt, U. Boehm, W. Meyerhof, N.I. Steinle, and S.D. Munger. 2015. TAS2R bitter taste receptors regulate thyroid function. FASEB Journal. 29:164–172.

Damak, S., M. Rong, K. Yasumatsu, Z. Kokrashvili, V. Varadarajan, S. Zou, P. Jiang, Y. Ninomiya, and R.F. Margolskee. 2003. Detection of sweet and umami taste in the absence of taste receptor T1r3. Science. 301:850–853.

Dazert, E., and M.N. Hall. 2011. mTOR signaling in disease. Current opinion in cell biology. 23:744–755.

Duan, P., C. Quan, W.T. Huang, and K.D. Yang. 2016. [PI3K-Akt/LKB1-AMPK-mTOR-p70S6K/4EBP1 signaling pathways participate in the regulation of testis development and spermatogenesis: An update]. Zhonghua nan ke xue = National journal of andrology. 22:1016–1020.

Foster, S.R., E.R. Porrello, B. Purdue, H.W. Chan, A. Voigt, S. Frenzel, R.D. Hannan, K.M. Moritz, D.G. Simmons, P. Molenaar, E. Roura, U. Boehm, W. Meyerhof, and W.G. Thomas. 2013. Expression, regulation and putative nutrient-sensing function of taste GPCRs in the heart. PloS one. 8:e64579.

Gentiluomo, M., L. Crifasi, A. Luddi, D. Locci, R. Barale, P. Piomboni, and D. Campa. 2017. Taste receptor polymorphisms and male infertility. Human reproduction. 32:2324–2331.

Gu, F., X. Liu, J. Liang, J. Chen, F. Chen, and F. Li. 2015. Bitter taste receptor mTas2r105 is expressed in small intestinal villus and crypts. Biochemical and Biophysical Research Communications.

Iwatsuki, K., M. Nomura, A. Shibata, R. Ichikawa, P.L. Enciso, L. Wang, R. Takayanagi, K. Torii, and H. Uneyama. 2010. Generation and characterization of T1R2-LacZ knock-in mouse. Biochemical and biophysical research communications. 402:495–499.

Kovacs, A., and R.H. Foote. 1992. Viability and acrosome staining of bull, boar and rabbit spermatozoa. Biotechnic & histochemistry : official publication of the Biological Stain Commission. 67:119–124.

Lee, E.C., D. Yu, J. Martinez de Velasco, L. Tessarollo, D.A. Swing, D.L. Court, N.A. Jenkins, and N.G. Copeland. 2001. A highly efficient Escherichia coli-based chromosome engineering system adapted for recombinogenic targeting and subcloning of BAC DNA. Genomics. 73:56–65.

Li, F. 2013. Taste perception: from the tongue to the testis. Molecular Human Reproduction.

Li, F., and M. Zhou. 2012. Depletion of bitter taste transduction leads to massive spermatid loss in transgenic mice. Mol Hum Reprod.

Li, N., and C.Y. Cheng. 2016. Mammalian target of rapamycin complex (mTOR) pathway modulates blood-testis barrier (BTB) function through F-actin organization and gap junction. Histology and histopathology. 31:961–968.

Liu, X., F. Gu, L. Jiang, F. Chen, and F. Li. 2015. Expression of bitter taste receptor Tas2r105 in mouse kidney. Biochemical and Biophysical Research Communications. 458:733–738.

Margolskee, R.F., J. Dyer, Z. Kokrashvili, K.S. Salmon, E. Ilegems, K. Daly, E.L. Maillet, Y. Ninomiya, B. Mosinger, and S.P. Shirazi-Beechey. 2007. T1R3 and gustducin in gut sense sugars to regulate expression of Na+-glucose cotransporter 1. Proceedings of the National Academy of Sciences of the United States of America. 104:15075–15080.

Masubuchi, Y., Y. Nakagawa, J. Ma, T. Sasaki, T. Kitamura, Y. Yamamoto, H. Kurose, I. Kojima, and H. Shibata. 2013. A novel regulatory function of sweet taste-sensing receptor in adipogenic differentiation of 3T3-L1 cells. PloS one. 8:e54500.

Max, M., Y.G. Shanker, L. Huang, M. Rong, Z. Liu, F. Campagne, H. Weinstein, S. Damak, and R.F. Margolskee. 2001. Tas1r3, encoding a new candidate taste receptor, is allelic to the sweet responsiveness locus Sac. Nature genetics. 28:58–63.

Meyer, D., A. Voigt, P. Widmayer, H. Borth, S. Huebner, A. Breit, S. Marschall, M.H. de Angelis, U. Boehm, W. Meyerhof, T. Gudermann, and I. Boekhoff. 2012. Expression of Tas1 taste receptors in mammalian spermatozoa: functional role of Tas1r1 in regulating basal Ca(2)(+) and cAMP concentrations in spermatozoa. PloS one. 7:e32354.

Mosinger, B., K.M. Redding, M.R. Parker, V. Yevshayeva, K.K. Yee, K. Dyomina, Y. Li, and R.F. Margolskee. 2013. Genetic loss or pharmacological blockade of testes-expressed taste genes causes male sterility. Proceedings of the National Academy of Sciences.

Nelson, G., J. Chandrashekar, M.A. Hoon, L. Feng, G. Zhao, N.J. Ryba, and C.S. Zuker. 2002. An amino-acid taste receptor. Nature. 416:199–202.

Nelson, G., M.A. Hoon, J. Chandrashekar, Y. Zhang, N.J. Ryba, and C.S. Zuker. 2001. Mammalian sweet taste receptors. Cell. 106:381–390.

Perez, C.A., L. Huang, M. Rong, J.A. Kozak, A.K. Preuss, H. Zhang, M. Max, and R.F. Margolskee. 2002. A transient receptor potential channel expressed in taste receptor cells. Nature Neuroscience. 5:1169–1176.

Ren, X., L. Zhou, R. Terwilliger, S.S. Newton, and I.E. de Araujo. 2009. Sweet taste signaling functions as a hypothalamic glucose sensor. Frontiers in integrative neuroscience. 3:12.

Somfai, T., S. Bodo, S. Nagy, E. Gocza, J. Ivancsics, and A. Kovacs. 2002. Simultaneous evaluation of viability and acrosome integrity of mouse spermatozoa using light microscopy. Biotechnic & histochemistry : official publication of the Biological Stain Commission. 77:117–120.

Sukcharoen, N., and J. Keith. 1996. Evaluation of the percentage of sperm motility at 24 h and sperm survival ratio for prediction of in vitro fertilization. Andrologia. 28:203–210.

Voigt, A., J. Bojahr, M. Narukawa, S. Hubner, U. Boehm, and W. Meyerhof. 2015. Transsynaptic Tracing from Taste Receptor Cells Reveals Local Taste Receptor Gene Expression in Gustatory Ganglia and Brain. Journal of Neuroscience. 35:9717–9729.

Voigt, A., S. Hubner, K. Lossow, I. Hermans-Borgmeyer, U. Boehm, and W. Meyerhof. 2012. Genetic labeling of tas1r1 and tas2r131 taste receptor cells in mice. Chemical Senses. 37:897–911.

Wauson, E.M., E. Zaganjor, A.Y. Lee, M.L. Guerra, A.B. Ghosh, A.L. Bookout, C.P. Chambers, A. Jivan, K. McGlynn, M.R. Hutchison, R.J. Deberardinis, and M.H. Cobb. 2012. The G protein-coupled taste receptor T1R1/T1R3 regulates mTORC1 and autophagy. Molecular cell. 47:851–862.

Wong, G.T., L. Ruiz-Avila, and R.F. Margolskee. 1999. Directing gene expression to gustducin-positive taste receptor cells. The Journal of neuroscience : the official journal of the Society for Neuroscience. 19:5802–5809.

Xu, J., J. Cao, N. Iguchi, D. Riethmacher, and L. Huang. 2012. Functional characterization of bitter-taste receptors expressed in mammalian testis. Molecular Human Reproduction.

Zoncu, R., A. Efeyan, and D.M. Sabatini. 2011. mTOR: from growth signal integration to cancer, diabetes and ageing. Nature reviews. Molecular cell biology. 12:21–35.

